# Hidden evolutionary constraints dictate the retention of coronavirus accessory genes

**DOI:** 10.1101/2023.10.12.561935

**Authors:** Stephen A. Goldstein, Teagan M. Feeley, Kristina M. Babler, Zoë A. Hilbert, Diane M. Downhour, Niema Moshiri, Nels C. Elde

## Abstract

Coronaviruses exhibit many mechanisms of genetic innovation, including the acquisition of accessory genes that originate by capture of cellular genes or through duplication of existing viral genes. Accessory genes influence viral host range and cellular tropism, but little is known about how selection acts on these variable regions of virus genomes. We used experimental evolution of mouse hepatitis virus (MHV) encoding a cellular AKAP7 phosphodiesterase and an inactive native phosphodiesterase, NS2 to model the evolutionary fate of accessory genes. After courses of serial infection, the gene encoding inactive NS2, ORF2, unexpectedly remained intact, suggesting it is under cryptic constraint uncoupled from the function of NS2. In contrast, AKAP7 was retained under strong selection but rapidly lost under relaxed selection. Experimental evolution also led to altered viral replication in a cell type-specific manner and changed the relative proportions of subgenomic viral RNA in plaque-purified viral isolates, revealing additional mechanisms of adaptation. Guided by the retention of ORF2 and similar patterns in related betacoronaviruses, we analyzed ORF8 of SARS-CoV-2, which arose via gene duplication and contains premature stop codons in several globally successful lineages. As with MHV ORF2, the coding-defective SARS-CoV-2 ORF8 gene remains largely intact, mirroring patterns observed during MHV experimental evolution, challenging assumptions on the dynamics of gene loss in virus genomes and extending these findings to viruses currently adapting to humans.

## Introduction

Coronaviruses have unusually large RNA genomes, ranging from ∼27-32 kilobases^1,2^, and incorporate a proofreading exonuclease into their RNA polymerase complex. Among RNA viruses, encoding a 3’-5’ exonuclease appears unique to coronaviruses and arenaviruses, and it is well-established that coronavirus exonucleases perform error-correction during infection^3–6^. Consequently, coronaviruses maintain larger genomes compared to most RNA viruses because the risk of catastrophic mutations is reduced. As a result, coronaviruses may be more tolerant to the addition of genetic material via mechanisms such as horizontal gene transfer^7–9^ and gene duplication more commonly associated with DNA viruses^10,11^. Intriguingly, a large 38.9 kb flavi-like virus sequence was recently assembled from a sea sponge but uses an alternative mechanism to maintain genome stability^12^.

Hypotheses on the evolutionary streamlining of RNA virus genomes^13–15^ predict that new genes, such as coronavirus accessory genes, are unlikely to persist in virus populations if they do not encode fitness benefits. Experimental studies of tobacco etch virus (TEV) support the assumption that exogenous genetic material may be rapidly purged under relaxed selection^14,16^. However, exogenous genes engineered into TEV, a small single-stranded positive sense RNA virus, substantially increase genome size, which might impart purifying selection that overwhelms any immediate fitness benefit afforded by the new gene product and drive its deletion. To date, no comparable studies using animal RNA viruses to test the interplay between gain of beneficial coding sequence and selection on genome size have been reported. Therefore, our understanding of how viral genome architecture and content evolve is limited, relative to the larger focus on how viruses evolve by nucleotide substitution. Given the potential for major evolutionary leaps afforded by gene rearrangements, gains, and losses this represents a fundamental gap in our understanding of viral evolution.

To explore how selection acts on newly acquired viral genes we used experimental evolution of mouse hepatitis virus (MHV), a prototypical betacoronavirus in the genus that includes SARS and MERS-related coronaviruses. ORF2 of MHV and related viruses encodes a 2’-5’ phosphodiesterase (PDE), NS2, that antagonizes the OAS-RNase L antiviral pathway. We used recombinant MHV encoding a cellular PDE, the AKAP7 central domain (AKAP7), as a functional replacement for an inactive NS2^17^ with a substitution at position 126 (His126Arg), that results in restricted viral replication^18^. Viral PDEs were derived via capture of cellular AKAP7-like genes^9,19^ so using this virus, MHV^AKAP7^, approximates a horizontal transfer event giving rise to a novel viral gene. Critically, the AKAP7 PDE is inserted in place of ORF4, using the native ORF4 transcriptional regulatory sequence (TRS) and surrounding sequence context^17^, allowing us to probe the relationship between selection and gene retention without substantial expansion of the genome or introduction of non-native regulatory sequence.

We found that the retention of coronavirus accessory gene sequences in different selective environments can in some instances override patterns of genome streamlining. Under relaxed selection in the absence of active OAS-RNase L immune pressure, AKAP7 was rapidly lost, consistent with the impact of selective constraints on genome size. However, in striking contrast, MHV ORF2 was retained under all conditions tested, despite encoding an inactive PDE. In parallel, we found that ORF8 of SARS-CoV-2 is largely intact even in lineages where it has lost coding capacity^20^. These results suggest widespread constraints on streamlining of viral genomes due to genetic information with hidden functions beyond protein coding capacity.

## Results

### Cell type-dependent selective pressure on viral phosphodiesterases

To test the impacts of distinct selective pressures on coronavirus genome composition, we used a recombinant mouse hepatitis virus (MHV) that has an inactivating H126R amino acid change in NS2, its native PDE, which is functionally complemented by insertion of the coding sequence for the cellular AKAP7 central domain PDE (AKAP7)^17^ (**Figure 1A-B**). The AKAP7 gene inserted into MHV is 618 nucleotides and replaces the 321 nucleotide ORF4, so expansion of the genome is minimal (∼1%). Additionally, MHV strain A59, the backbone for the viruses used in this study, has a fragmented ORF4 which produces a 2.2 kDa peptide and a theoretically novel ORF that is not translated^21^. Therefore, removal of ORF4 should not introduce any constraint due to loss of a functional protein. MHV with inactive NS2 (MHV^NS2Mut^) is restricted in mouse primary bone marrow-derived macrophages (BMDMs) by OAS-RNase L (ref^18^), while AKAP7 inserted into MHV (MHV^AKAP7^) compensates for the inactive NS2 and restores replication to wild-type virus levels in BMDMs and mouse liver. MHV^NS2mut^ is restricted in immortalized primary macrophages^22^, but replicates to wild-type levels in L2 mouse fibroblasts (**Figure 1C**), providing differential selection for courses of experimental evolution^18^. Similarly, MHV^AKAP7^ replicated to significantly higher titers than MHV encoding an inactive AKAP7 PDE (MHV^AKAP7mut^) (**Figure 1C**), demonstrating that these cells have an active OAS-RNase L pathway, which we hypothesized imposes selective pressure to retain an active PDE. In contrast, MHV^AKAP7^ and MHV^AKAP7mut^ replicated to an equivalent level in L2 fibroblasts, establishing these cells as a model for MHV^AKAP7^ evolution under relaxed selection (**Figure 1C**). Notably, the MHV^AKAP7^ viruses replicated to significantly lower titers than MHV viruses containing the native ORF4, suggesting some deleterious impact from the removal of ORF4, even though it does not encode a functional protein in MHV strain A59^21^.

**Figure 1.**
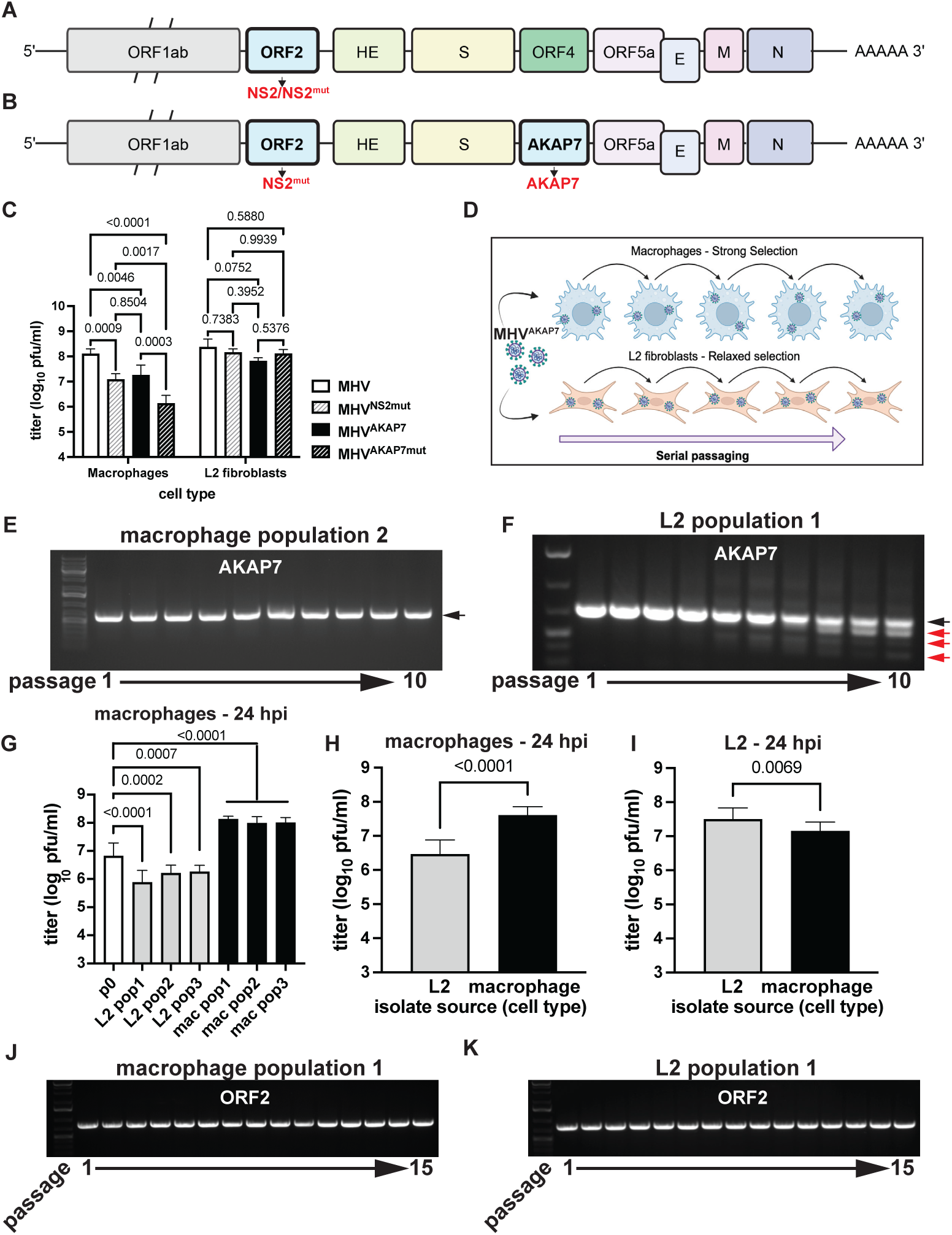
Experimental evolution of mouse hepatitis virus reveals hidden selective constraints. A) Schematic of the MHV genome with ORF2 and its PDE protein product NS2, highlighted. B) Schematic of the MHV^AKAP7^ genome, showing the inactive NS2 protein product and insertion of the AKAP7 gene in place of ORF4. C) Replication of wild-type MHV, MHV^NS2mut^, MHV^AKAP7^, and MHV^AKAP7mut^ in macrophages and L2 fibroblasts 24 hours post-infection at an MOI of 0.01. Statistical testing was conducted by 2-way ANOVA. One experiment representative of three is shown. D) Schematic of the experimental evolution protocol using conditions of strong and relaxed selection. E-F) PCR analysis of AKAP7 at passage 1 to 10 in macrophages and L2 fibroblasts, respectively. The black arrow indicates full-length intact AKAP7, while red arrows indicate AKAP7 amplicons with deletions. PCR analysis of AKAP7 in the remaining two macrophage and L2-passaged virus populations is available in **Figure S1A-D**. G) Replication in macrophages 24 hpi (MOI=0.01) of p0 MHV^AKAP7^ and p10 MHV^AKAP7^ populations from L2 fibroblasts and macrophages. Merged data from three independent experiments is shown and was subjected to statistical testing by 2-way ANOVA. H) Replication of plaque purified isolates in macrophages. Values are the average titer of all plaques. Statistical testing was done by unpaired t-test. I) Replication of plaque purified isolates in L2 fibroblasts. Values are the average titer of all plaques. Statistical testing was done by unpaired t-test. J-K) PCR analysis of ORF2 in macrophages and L2 fibroblasts, respectively. Images are representative of all three virus populations. Additional data related to this Figure can be found in **Figure S1.**

### Deletion of AKAP7 occurs rapidly under relaxed selection

Our experimental evolution workflow uses serial passage in macrophages and L2 cells to approximate evolution of a horizontally acquired PDE under different conditions (**Figure 1D**). We plaque purified passage 0 (p0) MHV^AKAP7^ and passaged this virus in triplicate in each cell type at a multiplicity of infection (MOI) of 0.01. At each passage we titrated virus replication by plaque assay to calculate the MOI for the subsequent passage and collected total RNA for analysis of AKAP7 by PCR and Sanger sequencing. In macrophages, full-length AKAP7 was retained through ten passages (**Figure 1E, Figure S1A-B**), whereas in L2 cells, smaller PCR amplicons appeared by passage three or four (**Figure 1F, Figure S1C-D**). The smaller bands became increasingly prominent through p10, while the full length AKAP7 band faded in intensity, indicating viruses with the full-length gene were disappearing from the population (**Figure 1F, Figure S1C-D**). We cloned and Sanger sequenced full-length bands from macrophages and L2 cells, and smaller bands representing putative AKAP7 degradation products from L2 cells at each passage.

AKAP7 from both cell types contained no point mutations compared to the sequence at p0 (raw data available on Figshare). The degradation bands that increased in frequency throughout passaging in L2 cells all contained AKAP7 sequence, but with large deletions (Figshare). The AKAP7 degradation products did not contain point mutations, presumably reflecting the high fidelity of coronavirus replication. This may suggest a lower barrier to genetic change via indels rather than nucleotide substitutions, which is consistent with findings characterizing the evolution of SARS-CoV-2^23^. Virus populations recovered at p10 from L2s exhibited significantly restricted replication relative to macrophage-passaged virus populations (**Figure 1G**), demonstrating that despite the heterogeneity in AKAP7 deletion patterns between replicates, all three MHV^AKAP7^ populations lost the ability to effectively suppress OAS-RNase L. Macrophage-passaged viruses replicated to significantly higher titers than at passage 0, suggesting adaptation to macrophages over time (**Figure 1G**)

We then plaque purified virus isolates from p10 virus populations from both cell types and used these isolates to infect macrophages and L2 cells. Consistent with the loss of AKAP7 during L2 passage, p10 plaque purified isolates from these cells exhibited reduced replication in macrophages (**Figure 1H**) and, unexpectedly, exhibited significantly increased replication in L2 fibroblasts relative to macrophage-derived isolates (**Figure 1I)**. The rapid loss of AKAP7 under relaxed selection is consistent with an evolutionary model wherein newly encoded viral genes must provide a near-immediate advantage to escape deletion.

### Inactive MHV ORF2 is retained under strong and relaxed selection

Like the loss of AKAP7 under relaxed selection, there is an *a priori* prediction that ORF2, which encodes an inactive PDE in MHV^AKAP7^, would also be rapidly lost from the MHV genome. ORF2, encoding the H126R inactivating substitution in NS2 should be dispensable in both macrophages and L2 fibroblasts. Surprisingly, PCR of ORF2 (**Figure 1J-K**) at each serial passage in macrophages and L2s, and sequencing of ORF2 after 10 passages, revealed no deletions, novel mutations, or reversion of the H126R substitution (Figshare). Given the unexpected retention of inactive ORF2, we conducted five additional passages in both cell types to provide additional opportunity for deletion. We sequenced the macrophage-passaged viruses again at p15 to confirm there was no reversion at amino acid position 126 that might suggest ORF2 retention in these cells, even out to passage 15, was due to OAS-RNase L-mediated selection (**Figure 1J-K; Figshare**). This unpredicted retention of ORF2 raised the possibility that protein-coding coronavirus genes might be regularly retained under constraints distinct from protein function.

To investigate whether similar patterns of constraint exist in the wild, we analyzed ORF2 from other betacoronaviruses in the same subgenus as MHV. Rabbit coronavirus HKU14^24^ has a premature stop codon mutation truncating NS2 to 43 amino acids and additional downstream stop codons in the same frame. However, the entire nucleotide sequence encoding full-length NS2 is present in four of five HKU14 sequences deposited in the National Center for Biotechnology Information databases, while the other has ∼100 bp of missing sequence (Figshare). In contrast, ORF2 in porcine hemagglutinating encephalomyelitis virus (PHEV) appears much more evolutionarily malleable^25–28^, with various isolates containing ORF2 with premature stop codons or deletions (Figshare). These findings are broadly consistent with constraints on deletion of ORF2, with the heterogeneity in PHEV suggesting retention of ORF2 is not as strict in some cases.

### Long read sequencing confirms retention of ORF2 after experimental evolution

To analyze ORF2 and AKAP7 genes more extensively after experimental evolution we performed Oxford Nanopore direct cDNA sequencing of plaque purified MHV^AKAP7^ p0, five p10 isolates from L2 cells, and one p10 isolate from macrophage infections. PCR analysis of the plaque purified isolates showed AKAP7 was uniformly intact in macrophage-derived isolates but deleted in L2- derived plaque isolates (**Figure S2A-B**). Mean read length across direct cDNA Nanopore sequencing runs ranged from 1,065 to 1,528 bases and average coverage was 2,603x to 27,296x per nucleotide of the MHV^AKAP7^ genome with a mean of 8,943x (**Table S1**). Across all sequencing runs ∼70-80% of reads aligned to the MHV^AKAP7^ genome (**Table S1**). To quantify changes in ORF2 and AKAP7 gene content after ten passages and control for variable overall sequencing depth across runs, we normalized average coverage of ORF2 and AKAP7 to average coverage of ORF1ab for each isolate. Consistent with PCR analysis, average relative coverage of ORF2 was the same in plaque purified passage 10 isolates from macrophages and L2 fibroblasts (**Figure 2A-C; Figure S2C-F**), demonstrating that this genomic region is under an unexpected constraint. The average relative AKAP7 coverage for p0 and macrophage p10 were 11.2 and 27.8, respectively, suggesting possibly enhanced expression of the viral AKAP7 gene (**Figure 2D-E**). In contrast, the average relative AKAP7 coverage for L2 p10 isolates was 0.332, a 33-fold decrease from p0, consistent with PCR analysis. (**Figure 2F; S2A-B; S2G-J)**. Among L2-derived isolates only isolate 4 retained some of the AKAP7 gene, specifically the short intergenic regions immediately 5’ and 3’ of the coding region (**Figure S2I**).

**Figure 2.**
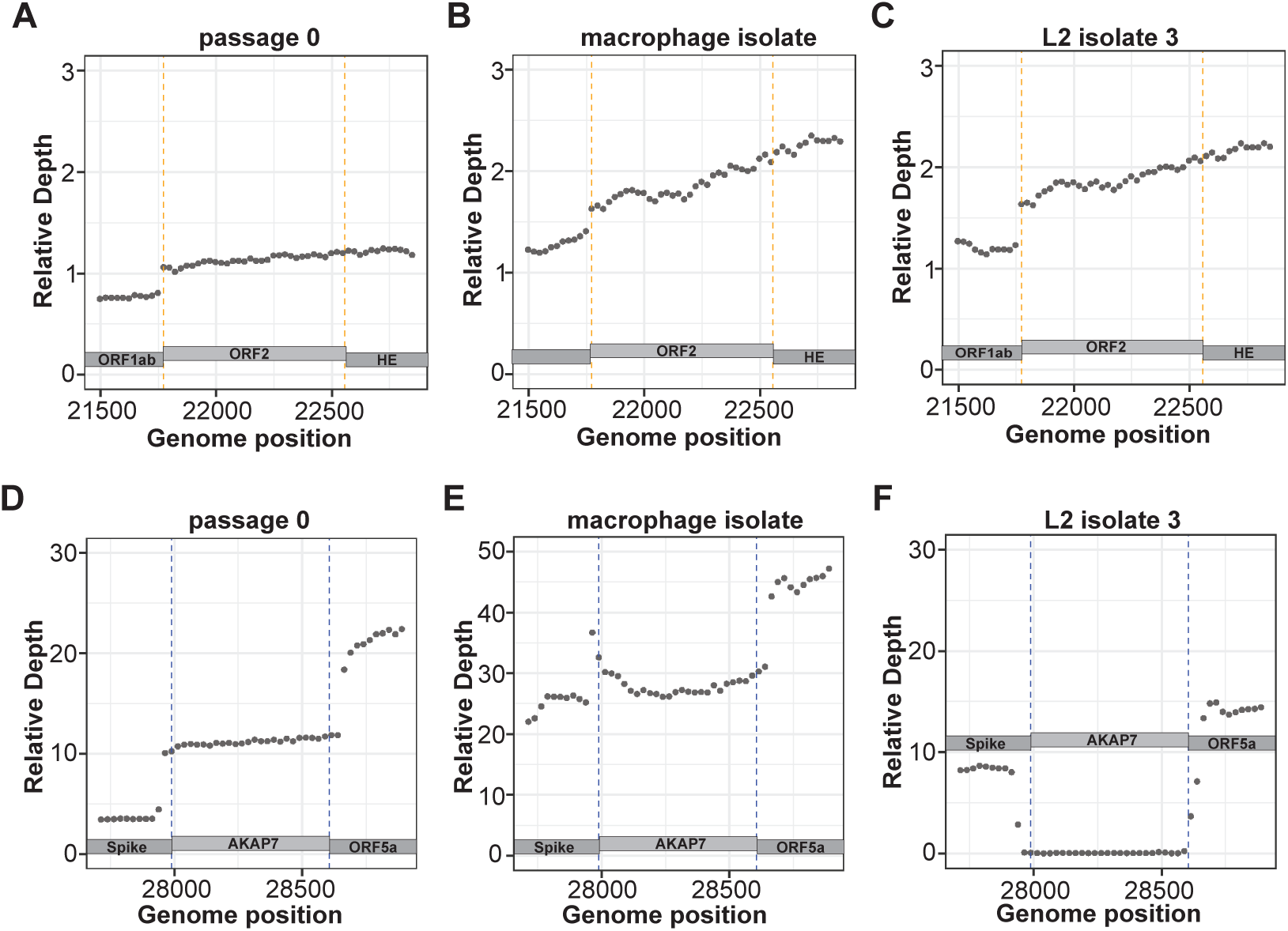
Nanopore direct cDNA sequencing shows retention of ORF2 and near-complete loss of AKAP7 following serial passage. A-C) Relative coverage depth plots of ORF2 in purified plaque isolates from passage 0 MHV^AKAP7^, and passage 10 macrophage and L2 fibroblast purified plaques. D-E) Relative coverage depth plots of AKAP7 in purified plaque isolates from passage 0 MHV^AKAP7^, and passage 10 macrophage and L2 fibroblast purified plaque isolates. PCR analysis of AKAP7 in plaque-purified virus isolates, as well as coverage plots from additional L2-derived isolates that were sequenced are available in **Figure S2**.

### Loss of AKAP7 increases expression of ORF2 and spike subgenomic RNA

In isolates where AKAP7 was lost during passaging in L2 cells, we used long read, direct cDNA sequencing data to examine changes in subgenomic (sg)RNA abundance with potential for functional impacts. Coronaviruses produce structural and accessory proteins through a nested set of sgRNAs, allowing for in-depth analysis of viral gene expression^29^. We identified subgenomic sgRNAs corresponding to each open reading frame by first filtering for reads containing unique leader-body TRS junctions previously identified for MHV strain A59 and filtered further to capture only those that contained the first 17 nucleotides of the expected gene^30^. We then calculated the abundance of each sgRNA as a percentage of the total canonical sgRNAs in the sample (Table S2).

Four of five L2-derived isolates exhibited a ∼3-to-5-fold increase in the relative abundance of ORF2 and S sgRNAs, the ORFs immediately upstream of the deleted AKAP7 (**Figure 3A-B**) and all five showed complete loss of the AKAP7 sgRNA (**Figure 3C**). Isolate 4, however, did not exhibit any change in the relative abundance of the ORF2 and S sgRNA (**Figure 3A-B**), corresponding with retention of the AKAP7 TRS, which is the native ORF4 TRS, immediately upstream as well as the more proximal 5’ and 3’ ends of the gene. Accordingly, isolate 4 exhibited use of the AKAP7 leader/TRS junction at similar relative frequency to passage 0 virus (**Figure 3D**), suggesting that loss of the AKAP7 TRS in the other isolates increased ORF2 and S sgRNA synthesis. The increased relative abundance of these sgRNAs is consistent with a mechanism where loss of the AKAP7 TRS facilitates more frequent synthesis of antisense ORF2 and S subgenomic RNAs, relative to the baseline frequency of other sgRNAs. Among all plaque-purified isolates, L2-passaged isolates replicated to significantly higher titers than macrophage-passaged isolates in L2 cells (Figure 1I). Among the representative isolates we sequenced, the relative increase in S sgRNA in four of them was associated with greater replication in L2 cells than the single macrophage-passaged isolate (**Figure 3H**), suggesting selection for greater relative spike sgRNA synthesis. The increase in spike sgRNA may result in greater production of spike protein, enhancing virus production in the absence of the RNA-sensing, antiviral OAS-RNase L pathway. Intriguingly, isolate 4 replicated similarly to the other four L2-derived isolates we sequenced (**Figure 3H**), despite no relative increase in S sgRNA expression. This is likely due to the relative increase in the ORF5a/E sgRNA associated with retention of the AKAP7 TRS (**Figure 3E**), which if coupled with increased production of E protein could enhance particle assembly^31,32^. We hypothesize that increased ORF5a/E sgRNA synthesis in isolate 4 is due to loss of the AKAP7 coding sequence that results in the orientation of two TRSs (the AKAP7 TRS and ORF5a/E TRS) immediately upstream of the gene. We observed no changes in the relative abundance of M or N sgRNAs under strong or relaxed selection (**Figure 3F-G**). The macrophage-derived isolate exhibited a two-fold increase in relative expression of the AKAP7 sgRNA (**Figure 3C**), suggesting its increased expression in macrophages (**Figure 3H**) aids in enhanced suppression of OAS-RNase L activation.

**Figure 3.**
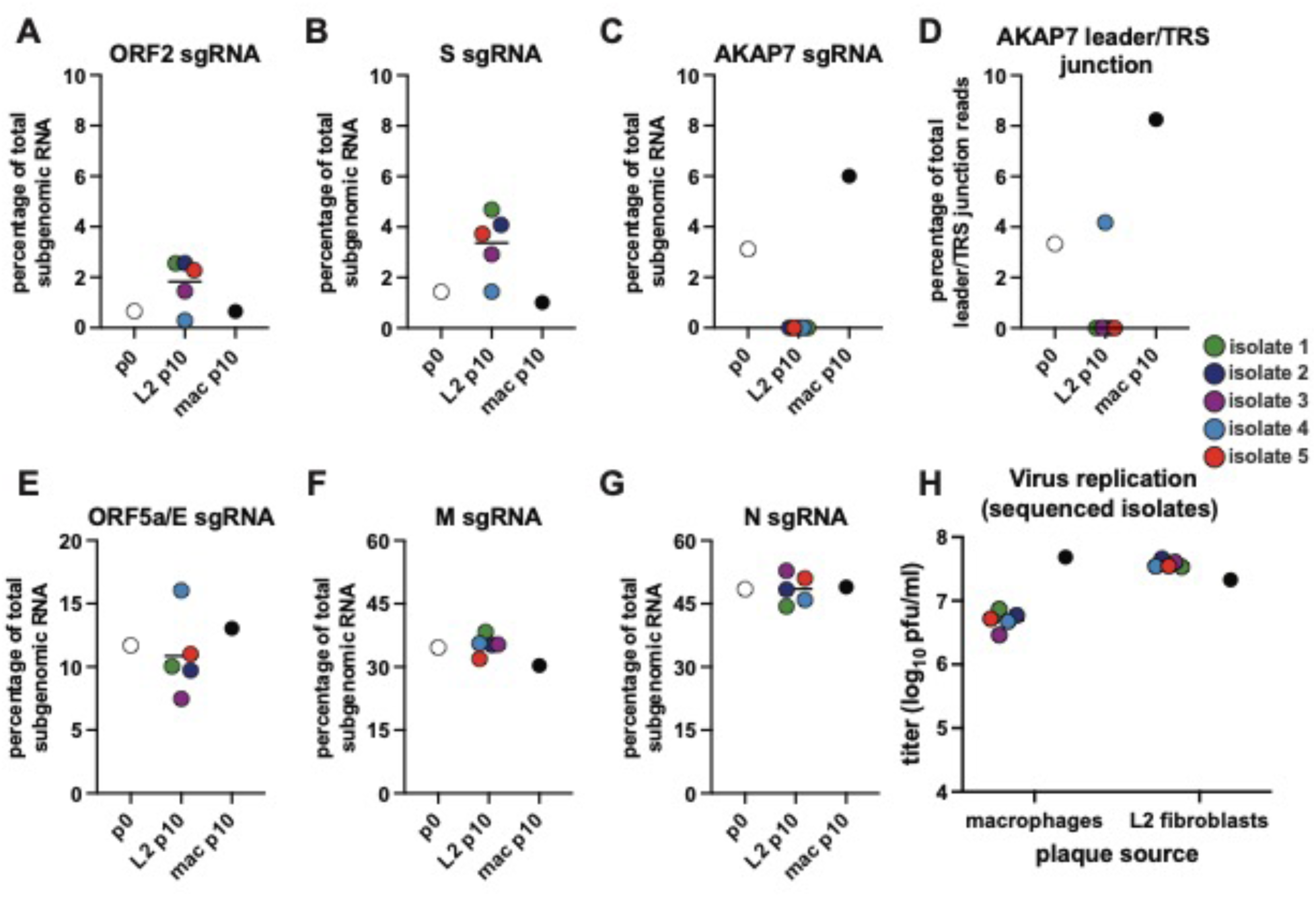
Experimental evolution of MHV^AKAP7^ alters relative expression of viral subgenomic RNA. A-C/E-G) Indicated sgRNAs plotted as a percentage of total subgenomic RNA for p0, L2 p10, and macrophage p10 passaged viruses. Individual L2 plaques are consistently colored across plots. D) Percentage of total reads with leader-body junctions that included the AKAP7-specific junction sequence. Run-statistics for all direct cDNA ONT sequencing are available in Table S1. Read-counts underlying the subgenomic RNA analysis are available in Table S2.

### SARS-CoV-2 ORF8 is retained despite loss of coding capacity

Returning to the persistence of ORF2 and related analysis in MHV-like viruses, we next tested whether similar constraints exist in other virus populations in the wild. The SARS-CoV-2 dataset in the Global Initiative on Sharing All Influenza Data (GISAID) database contains more than 16 million genomes, offering an unprecedented window into viral evolution. Analogous to the inactivation of MHV ORF2 in our system, SARS-CoV-2 ORF8 has acquired premature stop codons in numerous globally successful lineages, including B.1.1.7, XBB.1, XBB.1.5, XBB.1.9, and XBB.1.16^20^, and a mutation in the ORF8 transcriptional regulatory sequence (TRS) in the BA.5 lineage that substantially reduces ORF8 subgenomic RNA and protein synthesis (**Figure 4A-B**)^33^. ORF8 has been of interest since early in the SARS-CoV-2 pandemic due to a small deletion that emerged in its SARS-CoV ortholog in 2003^34^, an early cluster of infections in Singapore involving an ORF8 deletion variant^35,36^, as well as debate over its role during SARS-CoV-2 infection^10,36–43^. Despite the intense focus on ORF8 function and loss of full-length protein synthesis, very little attention has been devoted to the evolution of the gene itself.

**Figure 4.**
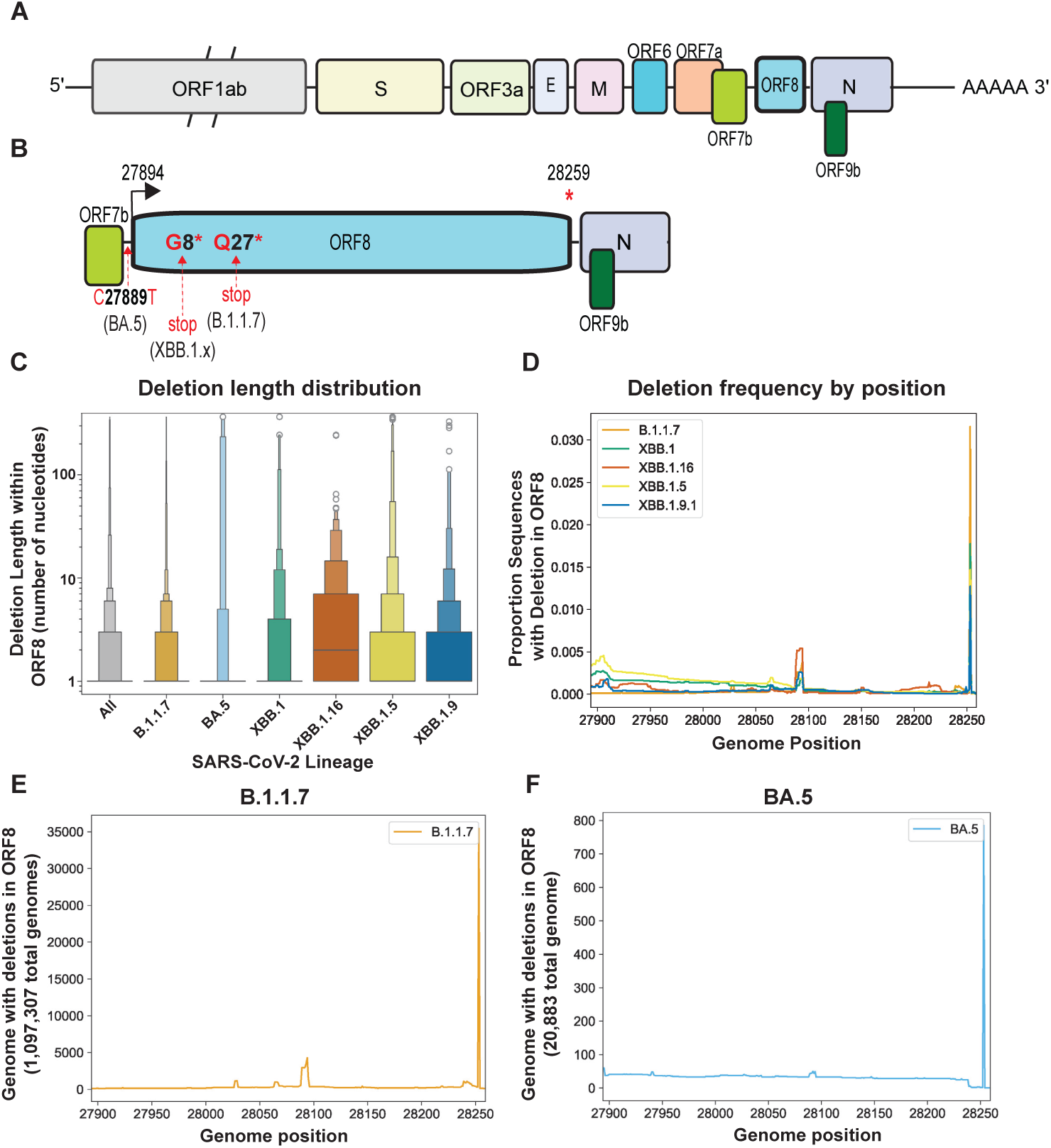
ORF8 is retained in SARS-CoV-2 lineages despite mutations that disrupt protein coding. A) Schematic of SARS-CoV-2 genome. B) Schematic (with ORF8 expanded, not to scale) of ORF8, with mutations resulting in loss of subgenomic mRNA synthesis (BA.5) or an early stop codon (B.1.1.7 and XBB.1.x) indicated. **Figure S3** contains the histograms showing the epidemic curve of each lineage. C) Deletion length distribution - this displays the size of all deletions in ORF8 in each lineage from 0 to >100 nucleotides. **Table S3** contains the raw data underlying this plot. D-F) Proportion of genomes from lineages with a deletion at each position in SARS-CoV-2 ORF8. **Figure S4** contains these plots for the XBB lineages we analyzed.

We downloaded all sequences assigned to the B.1.1.7, BA.5, XBB.1, XBB.1.5, XBB.1.9, and XBB.1.16 lineages from GISAID and applied a cutoff date to exclude any samples without metadata or with metadata suggesting they are incorrectly assigned (**Figure S3**). For each lineage, we further defined a start-date for circulation and calculated the date at which 50% and 90% of sequences assigned to this lineage were collected, as of November 29, 2023. We further excluded any sequences with any ambiguous nucleotides in the ORF8 gene to eliminate confounding of our analysis by poor-quality sequences, which might obscure *bona fide* deletions. We calculated a deletion length distribution of ORF8 in B.1.1.7 (1,097,307 sequences), BA.5 (20,833 sequences), XBB.1 (22,031 sequences), XBB.1.5 (166,068 sequences), XBB.1.9.1 (19.990 sequences), and XBB.1.16 (19,119 sequences) (**Figure 4C**). Across all lineages 96% of deletions were smaller than ten nucleotides, ranging from a high of 98.15% of deletions smaller than ten nucleotides for B.1.1.7 to a low of 86.6% of deletions smaller than ten nucleotides for XBB.1.16. (**Figure 4C, Supplementary File 5**). We then analyzed all sequences in the dataset to determine whether any regions of ORF8 were particularly deletion-tolerant or resistant. While XBB.1.5 ORF8 deletions were enriched in the 5’ half of the gene (**Figure 4D, S4B)**, deletions were uniformly rare at all positions across the other lineages we analyzed (**Figure 4D-F, S4A-D)** except for a single site at the 3’ end that we speculate may impact sgRNA synthesis of downstream ORFs N and 9b.

To determine whether ORF8 in these lineages degraded over time, we calculated the percent of ORF8 gene content in these lineages, sorted by collection date, and plotted these values for all genomes of all six lineages (**Figure 5A-F**). To determine how the appearance of ORF8 deletions in these lineages related to their epidemiological rise and fall, we calculated the dates at which 50% (dashed vertical line) and 90% (solid vertical line) of sequences assigned to each lineage had been collected. For all lineages, genomes with deletions were rare outside this window, generally occurring early after the appearance of a lineage and going extinct during lineage growth or emerging very late when the lineage was already in steep decline.

**Figure 5.**
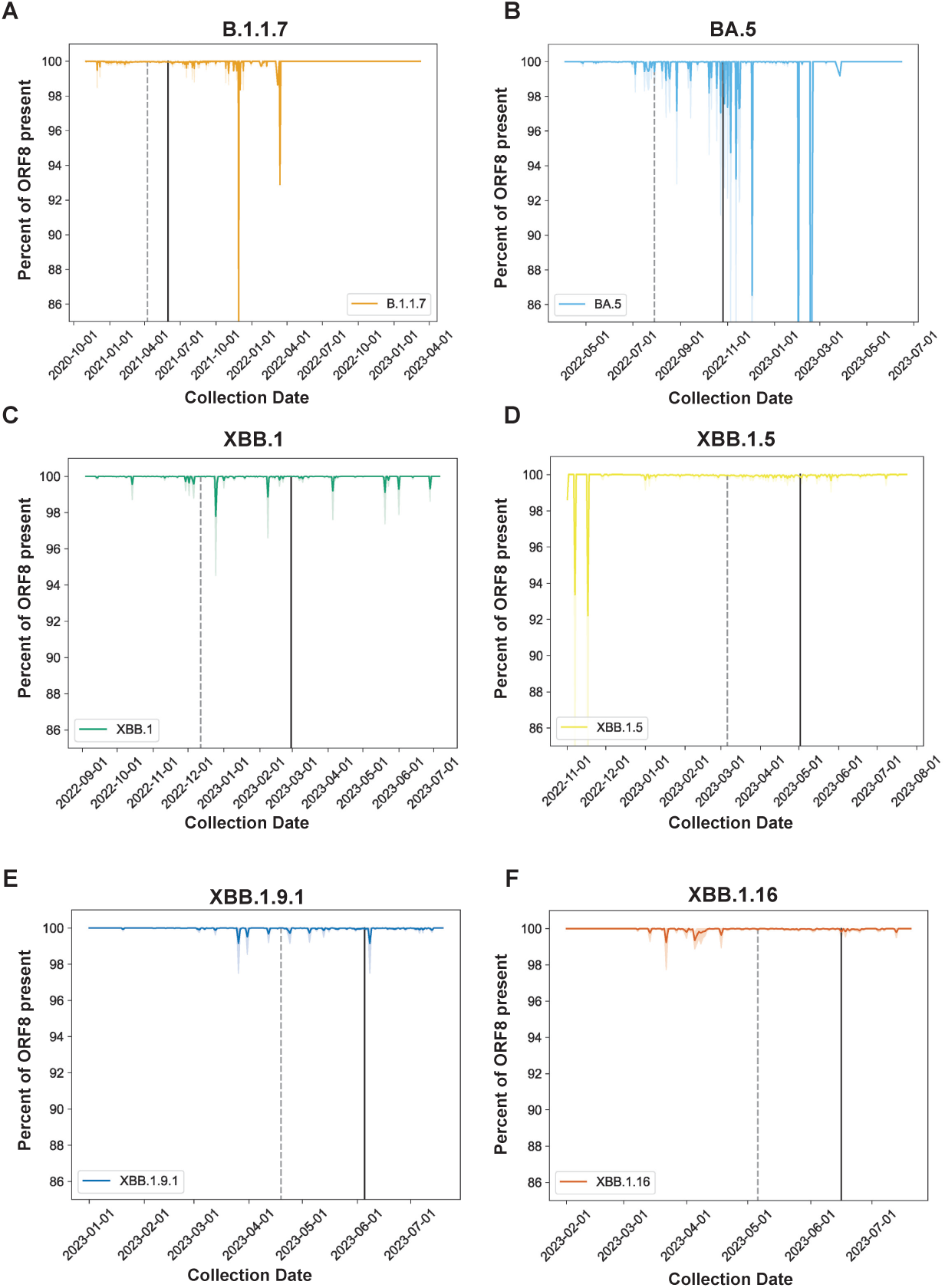
SARS-CoV-2 ORF8 is recurrently retained following acquisition of an early stop codon. A-F) Line plots of percent nucleotide content for the indicated SARS-CoV-2 lineages over time throughout the course of their circulation in humans. Dark lines are the mean percentage for all sequences collected on that date, and faded lines are the 95% confidence interval. Dashed vertical line is the date by which 50% of sequences assigned to the designated lineage were collected, and the solid line is the 90% cutoff. Figure S3 contains the histograms showing the epidemic curve of each lineage analyzed. Figure S4 contains plots of deletions at each position in ORF8 for the XBB lineages we analyzed.

## Discussion

Findings from this study add to a growing recognition that the evolutionary fate of protein coding regions do not hinge on protein function alone. In this case, the sustained, unexpected retention of ORF2 in both macrophages and L2 fibroblasts through fifteen passages suggests a complex interplay of selective pressure acting on viral coding sequence. In L2s the PDE activity of ORF2 is dispensable and the stability of the H126R substitution during experimental evolution in macrophages is consistent with AKAP7 fully counteracting OAS-RNase L activity. While it is formally possible that an unknown secondary role for the ORF2-encoded protein NS2 antagonizes innate immunity, no such function has been identified and the protein contains no identifiable domains other than the PDE.

Analyzing the relative abundance of subgenomic RNAs helped bring into focus the potential impact of genomic changes on viral replication under experimental selection. Increases in spike sgRNA associated with loss of AKAP7 and its transcriptional regulatory sequence (TRS) suggests that serial passage in L2 cells provides for multiple adaptive pathways, not solely selection on genome size. The loss of AKAP7 suggests this exogenous gene lacks any major regulatory function in the context of the MHV genome, while the associated increase in spike sgRNA synthesis suggests an adaptive increase in infectivity that is available in the absence of pressure from OAS-RNase L. Increased ORF5a/E but not spike sgRNA synthesis in one of the other L2-derived isolates revealed another pathway even over short courses of experimental evolution. These changes impacting synthesis of an adjacent sgRNA also suggest that retention of ORF2 may be influenced by similar forces. Increased AKAP7 sgRNA synthesis correlated with increased replication in macrophages, experimentally demonstrating that alterations in viral gene expression can favor virus replication in ways distinct from amino acid substitutions that alter viral protein function.

Under a simple model of selection favoring genome streamlining, viruses with ORF2 or ORF8 deleted would quickly dominate populations soon after the potential to encode full-length protein was lost. The exact nature of the evolutionary constraint on MHV ORF2 and SARS-CoV-2 ORF8 is not yet clear. A leading possibility is that at least some coronavirus genes contain regulatory functions in addition to their protein coding capacity that tune the transcription of other genes. MHV ORF2 is upstream of spike, albeit with the defective HE gene intervening, and SARS-CoV-2 ORF8 immediately precedes nucleocapsid, which has multiple functions key to promoting viral fitness^44^. If accessory genes promote transcription of spike, nucleocapsid, and other subgenomic RNAs, the loss of such genes might impact viral fitness, and lineages containing such deletions would go extinct. Results from experimental evolution of MHV are consistent with this idea given the link between genomic deletions and altered expression of proximal viral genes. Increased transcription of subgenomic RNAs encoding innate immune antagonists has been linked to enhanced suppression of innate immunity in major SARS-CoV-2 variants of concern^45^, which is consistent with our hypothesis and analysis of SARS-CoV-2 ORF8 retention.

Similar observations are evident from experimental deletion of accessory genes in SARS-CoV-2. For example, deletion of the ORF6 coding region impacts sgRNA levels of spike, nucleocapsid and the accessory protein ORF3a (^46^). Additionally, increased transcription of multiple sgRNAs encoding innate immune antagonists (ORF9b, ORF6, N and N*) has been linked to enhanced suppression of the innate immune system in major SARS-CoV-2 variants of concern^33,45^. Although two directly adjacent TRSs have not been observed in SARS-CoV-2, analogous to what we identified in L2-passaged isolate 4, increased expression of SARS-CoV-2 ORF9b has been linked to a triple nucleotide change that strengthens homology between the ORF9b TRS and the leader TRS^33^. Continued surveillance and analysis of SARS-CoV-2 evolution is key to revealing the intricate relationship between changes in other sgRNAs and the selective pressures acting on ORF8 and its regulatory regions.

Exceptions to these patterns of constraint include the loss of ORF2 in HKU1 viruses and the collection of some SARS-CoV-2 genomes with rare ORF8 deletions. In the case of SARS-CoV-2, any decreased fitness resulting from ORF8 deletion might be balanced by adaptations that compensate for loss of ORF8, affording fitness benefits of genome compression while shedding the deleterious impact of losing the gene. Continued surveillance of SARS-CoV-2 might reveal such variants and provide more insight into fundamental factors influencing coronavirus evolution.

Identifying additional patterns of evolutionary constraint may reveal other sequence features impacting viral fitness. Patterns of selection can be prioritized for mechanistic studies using reverse genetics systems established for MHV and SARS-CoV-2, opening a bridge between experimentation and computational analysis of viral evolution. Experimentally dissecting fitness impacts will benefit from future competition studies. For example, the loss of ORF2 in HKU1 results in viable viruses and fitness defects of MHV^ΔORF2^ might appear only in direct competition with wild-type viruses and not in parallel courses of experimental evolution. Overall, this work demonstrates the power of merging experimental approaches with computational analysis of massive datasets to identify patterns of constraint influencing the evolution of emerging and pandemic viruses.

## Resource Availability

### Lead contact

Requests for further information and resources should be directed to and will be fulfilled by the lead contact, Nels Elde.

### Materials Availability

- This study did not generate new or unique reagents
- Viruses used in this study were obtained from Dr. Susan Weiss at the University of Pennsylvania Perelman School of Medicine. Requests for these viruses can be fulfilled by the lead contact, Dr. Nels Elde, with permission from Dr. Susan Weiss or by Dr. Weiss directly.
- L2 and 17-Clone 1 cells were obtained from Dr. Susan Weiss at the University of Pennsylvania Perelman School of Medicine. Requests for these viruses can be fulfilled by the lead contact, Dr. Nels Elde, with permission from Dr. Susan Weiss or by Dr. Weiss directly.
- Macrophages were obtained from Dr. Sunny Shin at the University of Pennsylvania Perelman School of Medicine, Department of Microbiology. Requests for these cells can be fulfilled by the lead contact, Dr. Nels Elde, with permission from Dr. Sunny Shin or by Dr. Shin directly.
- All other reagents are available from the lead contact, Dr. Nels Elde without restriction.

### Data and Code Availability

#### Data

- All data (fasta files) from Sanger Sequencing of AKAP7 and ORF2 genes are available at Figshare (Figshare: 10.6084/m9.figshare.26937838, 10.6084/m9.figshare.26937637)
- The alignment of HKU14, and HKU1, and OC43 as well as the PHEV ORF2 alignment are available at Figshare (Figshare: 10.6084/m9.figshare.26939662)
- Raw sequencing data (fastq files) from Oxford Nanopore sequencing are available at NCBI under BioProject ID PRJNA1031673

#### Code

- All code used to analyze SARS-CoV-2 genomes is available on GitHub at https://github.com/niemasd/SC2-Deletion-Analysis
- Requests for additional information concerning analysis of Oxford Nanopore sequencing are available by request from the lead contact Dr. Nels Elde without restriction.

## Acknowledgements

This work was supported by NIH grants F32AI152341 to SAG and R35GM134936 to NCE. We thank Susan Weiss and Sunny Shin at the University of Pennsylvania for reagents. We thank Lucy Thorne (Imperial College of London) and Tom Peacock (Pirbright Institute), as well as Harriet Mears and David Bauer from the Francis Crick Institute for helpful discussions.

## Author Contributions

**SAG** and **NCE** conceived the project, analyzed data, designed figures and wrote, edited, and revised the manuscript. **SAG** was principally responsible for the design and oversight of all experiments, measured viral titers at the onset of the project and throughout, and conducted polyA selection of total RNA for direct cDNA sequencing. **TMF** conducted serial passaging experiments and associated cloning, Sanger sequencing sample prep, and viral titration. **KMB** conducted serial passaging as well as cloning and Sanger sequencing sample prep. **KMB** also was responsible for polyA selection of total RNA for direct RNA sequencing. **ZAH** analyzed direct cDNA sequencing coverage of the AKAP7 and ORF2 genes in experimentally evolved viruses. **DMD** prepared libraries for direct cDNA and direct RNA sequencing and conducted the subgenomic RNA analysis. **NM** conducted all analysis of SARS-CoV-2 genomes.

## STAR Methods

### Experimental Model and Study Participant Details

#### Cells and Viruses

Immortalized primary macrophages were provided by Sunny Shin^22^, 17-Clone 1 and L2 cells were provided by Susan Weiss. Macrophages were cultured in RPMI-1640 supplemented with 10% FBS, 1% L-glutamine, and 1% Penicillin/Streptomycin. 17-Clone 1 and L2 cells were cultured in DMEM supplemented with 10% FBS, 1% L-glutamine, and 1% Penicillin/Streptomycin. All cell lines were maintained at 37° C + 5% CO2. The cell lines were not authenticated, as none are commercially available, but exhibited expected morphologies and culture properties. The sex of the organism of origin of these cell lines is not known.

All viruses used in this study were provided by Susan R. Weiss at the University of Pennsylvania. Wild-type MHV strain A59 and MHV^NS2mut^ were previously described^18^ as were MHV^AKAP7^ and MHV^AKAP7mut17^. Viruses were received as seed aliquots and 10 µl of each virus was added to one T75 flask containing a confluent monolayer of 17 Clone-1 cells in 2 ml of DMEM+2% FBS. Flasks were incubated for one hour at 37° C, after which 10 ml of fresh DMEM+2% FBS was added, for a total volume of 12 ml. Once widespread cytopathic effect was present (18-24 hours post-infection) the flasks were put through three freeze-thaw cycles and the supernatant was clarified via centrifugation and aliquoted for later use. Viral stocks were titrated by plaque assay, as described below.

### Methods Details

#### Virus infections

One day after seeding in 12-well plates, macrophages or L2 cells were infected at a multiplicity of infection (MOI) of 0.01 in 200 µl volume and incubated for one hour at 37 ° C, with rocking every 15 minutes. After one hour, cells were washed 3 times with PBS and 1 ml of fresh media (RPMI or DMEM+2% FBS for macrophages and L2 cells, respectively) was added to each well. 24 hours post-infection the supernatant was collected and stored at -80° C. We also harvested total RNA for additional analyses (described below). Initial infections, all serial passages, and endpoint replication comparisons were done at an MOI of 0.01. Viral titers were quantified by plaque assay on confluent L2 cell monolayers. Experiments comparing replication of virus populations or plaque purified isolates were conducted in triplicate and replicated three times.

#### Plaque assays

Supernatants collected 24 hours post-infection were serially diluted 1:10 to an endpoint of 10^-8^ ^i^ DMEM+2% FBS, and the 10^-3^ to 10^-8^ dilutions were used to infect confluent L2 monolayers in 200 µl volume. Infected cells were incubated for 1 hour at 37 ° C with rocking every 15 minutes. After one hour 3 mL of semi-solid agarose overlay (1:3; 2% agarose:DMEM+2%FBS) was added to each well and the plates returned to 37 ° C for 36 hours. 36 hours post-infection the agarose overlay was removed and the monolayers were washed with PBS and stained with 0.5% crystal violet plus 20% methanol. The stain was washed off with water and plaques counted and recorded for analysis.

#### RNA extraction and cDNA synthesis

Total RNA was harvested 24 hours post-infection on each serial passage with the Zymo Research Quick-RNA Miniprep Kit (Cat. #R1057) following the manufacturer’s protocol. To increase yield we eluted the RNA in a reduced volume of 25 µl nuclease-free water. RNA quantification and quality assessment was conducted using a Synergy HT BioTek plate reader. cDNA was synthesized using the Thermo Scientific Maxima First Strand cDNA Synthesis Kit with dsDNase (Lot. 2664799) with 1 µg of input RNA.

#### PCR and Sanger sequencing

At each passage the AKAP7 central domain and ORF2 genes from MHV^AKAP7^ were analyzed by PCR. We used Phusion Flash HiFi Master Mix (Cat # F548S) in 20 µl/reaction with 2 µl of cDNA template. Thirty cycles of PCR were performed. Cycling conditions were: **1)** Denaturation, 98°C for 10 seconds **2) a)** Denaturation (98°C for 2 seconds) **b)** Annealing (63°C for 10 seconds) **c)** Extension (72°C for 30 seconds) **d)** Final extension (72°C for 2 minutes) **3)** Hold (4°C).

PCR products were visualized by agarose gel (1%) electrophoresis then extracted and purified using Zymoclean Gel DNA Recovery Kit (Zymo Research Lot. #213587) following manufacturer protocols. DNA was eluted in 10 µl. Purified PCR products were prepared for Sanger Sequencing by TOPO cloning using the Invitrogen TA Cloning™ Kit, with pCR™2.1 Vector (Cat. #K202020) following manufacturer protocols. Plasmid DNA was extracted using the Zippy Plasmid Miniprep Kit (Zymo Research Cat. # 11-308) following manufacturer protocol with elution 75 µl nuclease-free water. Plasmid DNA was quantified with a Synergy HT BioTek plate reader and sent for sequencing to the University of Utah DNA sequencing core or Genewiz.

#### Sequence analysis

Sequence files were imported into Geneious and aligned to either the AKAP7 PDE or ORF2 coding sequences using the MAFFT Geneious plug-in^47^. HKU1, HKU14, and PHEV multiple sequence alignments were likewise generated using the MAFFT Geneious plug-in with default parameters.

#### Oxford Nanopore direct cDNA sequencing

1 mL of media containing a plaque isolate was used to infect 2x 10 cm dishes of desired cell type (macrophages or L2). 18-20 hours post-infection total RNA was collected using Zymo Research Quick-RNA Miniprep Kit (Cat. #R1057). The same lysis buffer (1 ml) was used on both dishes/per infection to maximize yield. Polyadenylated RNA was extracted using the Invitrogen Poly(A)Purist MAG Kit (Cat # AM1922) following the manufacturer protocol. 100 ng PolyA+ RNA was prepared for sequencing using the Direct cDNA Sequencing Kit SQK-DCS109 from Oxford Nanopore Technologies, according to the manufacturer’s instructions. The prepared cDNA libraries were sequenced using the MinION Mk1C with FLO-MIN106D flow cells, version 9.4.1. Fastq files were generated from fast5 files using the high-accuracy model of Guppy basecaller, version 6.5.7 with the following parameters hac_guppy -c dna_r9.4.1_450bps_hac.cfg -x auto --compress_fastq.

#### Direct cDNA sequencing coverage analysis

Fastq files from Nanopore sequencing runs were individually aligned to the MHV^AKAP7^ (passage 0) reference genome. Alignments were performed using minimap2 (v.2.23) and subsequently sorted and indexed with samtools (v.1.16). Coverage at every position across the genome for each sample was calculated using the bedtools (v.2.26.0) genomecov command. Coverage was then normalized for each sample to the average coverage across ORF1ab (nucleotide positions 211-21746 in the reference genome), which contains the non-amplified reading frame, allowing for comparison across sequencing runs and samples. A rolling average of this normalized read depth was calculated across the genome in 25 bp windows and plotted with ggplot2 in R. For AKAP7 and ORF2 analysis, rolling averages were recalculated for the region containing the gene sequence ± 300 bp of flanking sequence on either side and plotted identically to the genome wide coverage plots. Fastq sequencing files are available in the Sequence Read Archive at BioProject ID PRJNA1031673.

### Subgenomic RNA analysis

Subgenomic RNAs were analyzed by concatenating fastq files of a given sample followed by searching for the subgenomic RNA of interest using the seqkit grep function. The subgenomic RNAs were initially identified by gene-specific leader/body junction sequences previously described for MHV strain A59^30^, and further refined by searching for the first 17 base pairs of a given ORF. This final refinement ensured that we captured only the canonical sgRNA associated with a given leader/body junction.

#### SARS-CoV-2 ORF8 Deletion Analysis

Lineage-specific SARS-CoV-2 sequence datasets were downloaded from GISAID on 2023-07-28. They were then aligned to the NC_045512.2 reference genome using ViralMSA v1.1.31 using its default settings, wrapping around Minimap2 v2.24. The sample collection date distributions, deletion length distributions, per-position non-deletion nucleotide counts, and per-position deletion counts were calculated and visualized for each lineage using an in-house Python script, which can be found with annotations in the Jupyter notebook within the following GitHub repository: https://github.com/niemasd/SC2-Deletion-Analysis/blob/main/Analysis.ipynb

### Statistical Analysis

Statistical testing in this study was conducted on comparisons of viral replication between virus populations or plaque purified isolates. Data analysis and testing was conducted in GraphPad Prism on log-transformed viral titers. Viral titers in Figure 1C and 1G were subjected to multiple comparison testing by 2-way ANOVA while viral titers in Figure 1H and 1I were compared by unpaired t-test. All comparisons were made using triplicate values as described in the Figure 1 legend.

### Key resources table

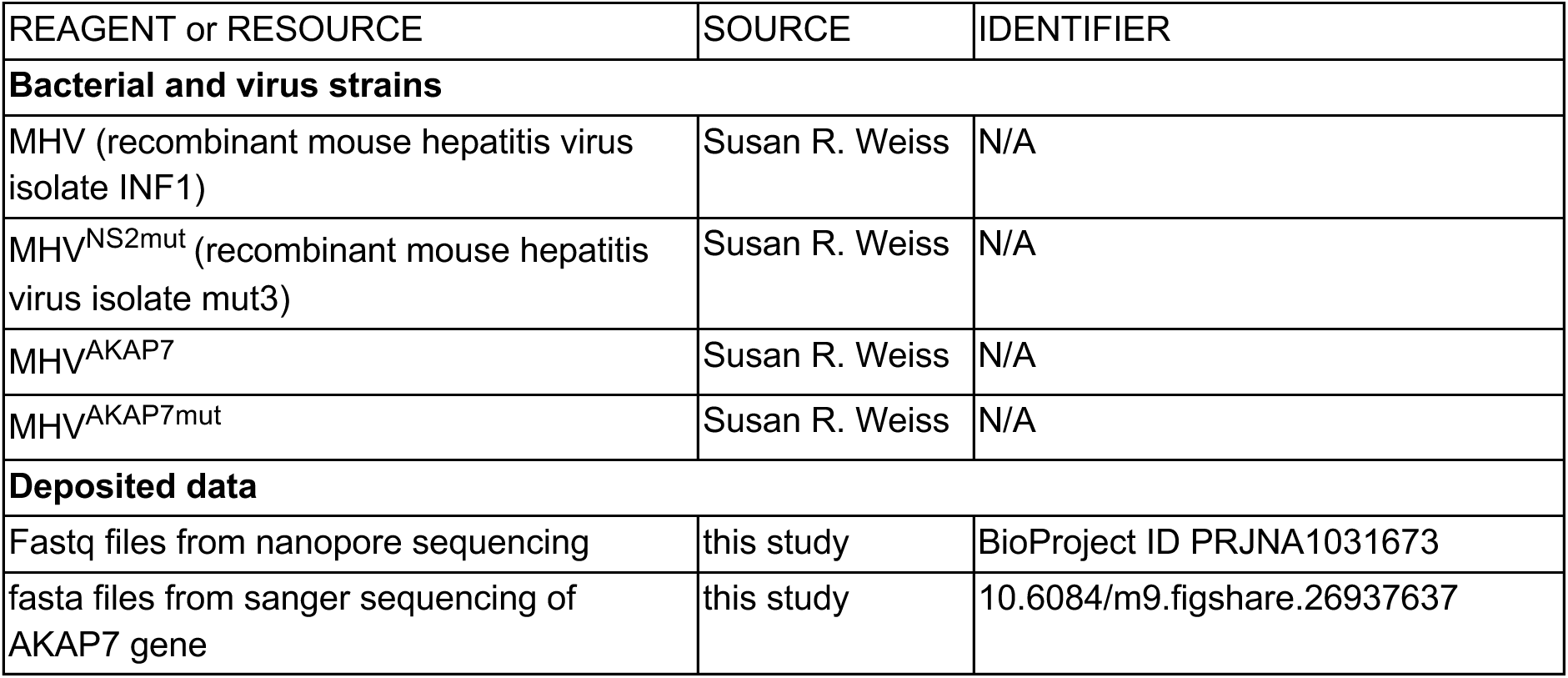
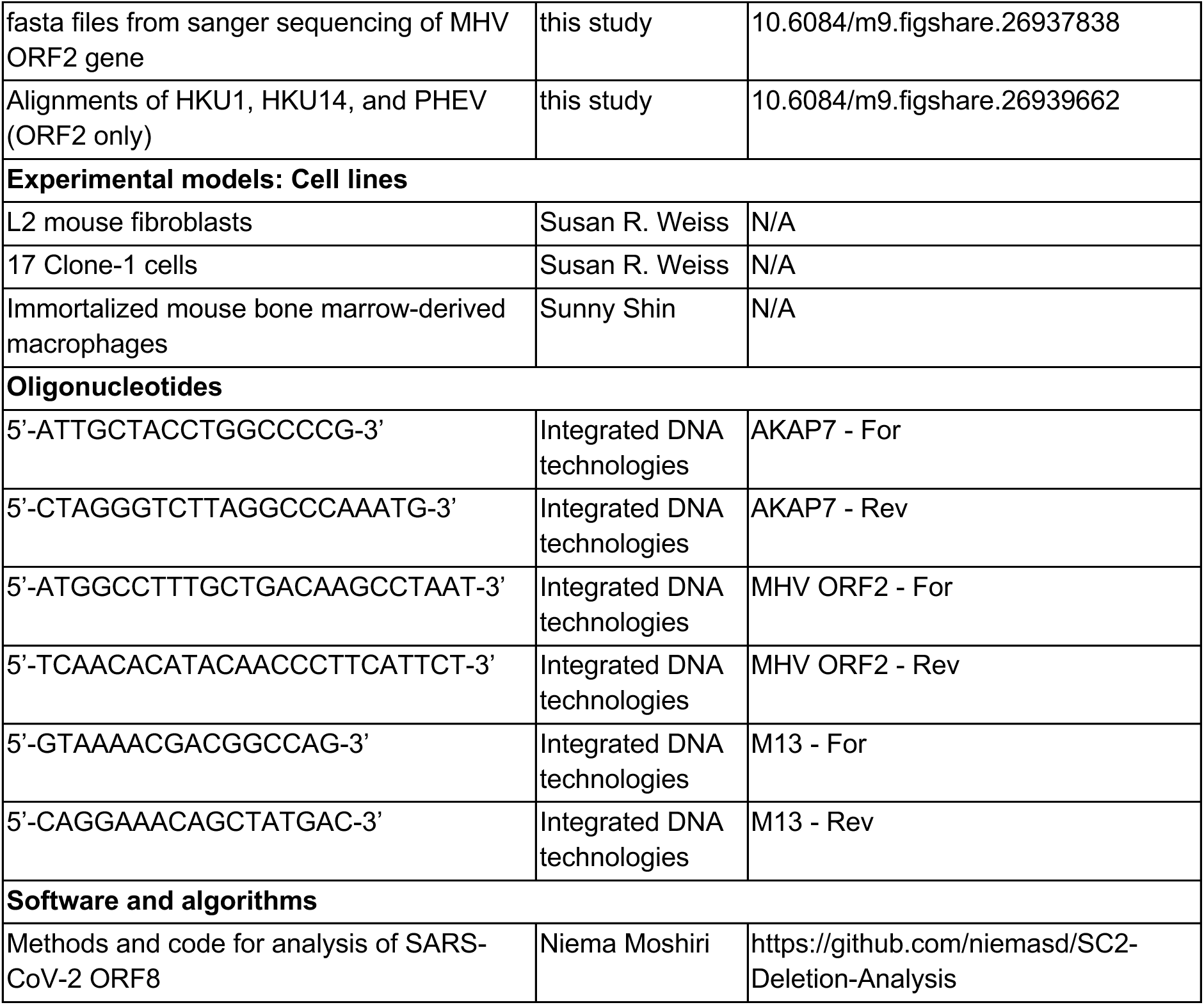

## Supplemental Table Legends

**Table S1. Oxford nanopore direct cDNA sequencing statistics, related to Figures 2 and 3.** This table reports data on coverage and read length for direct cDNA sequencing of MHV^AKAP7^ passage 0 and passage 10 viruses.

**Table S2. Viral subgenomic RNA counts and relative abundance raw numbers, related to Figure 3.** This table contains the counts of viral subgenomic RNA calculated using two different methods, as described. It also reports the relative abundance of each sgRNA as a percentage of total subgenomic RNA in the sample. This data underlies the plots in **Figure 3A-G**.

---

**Figure S1.**
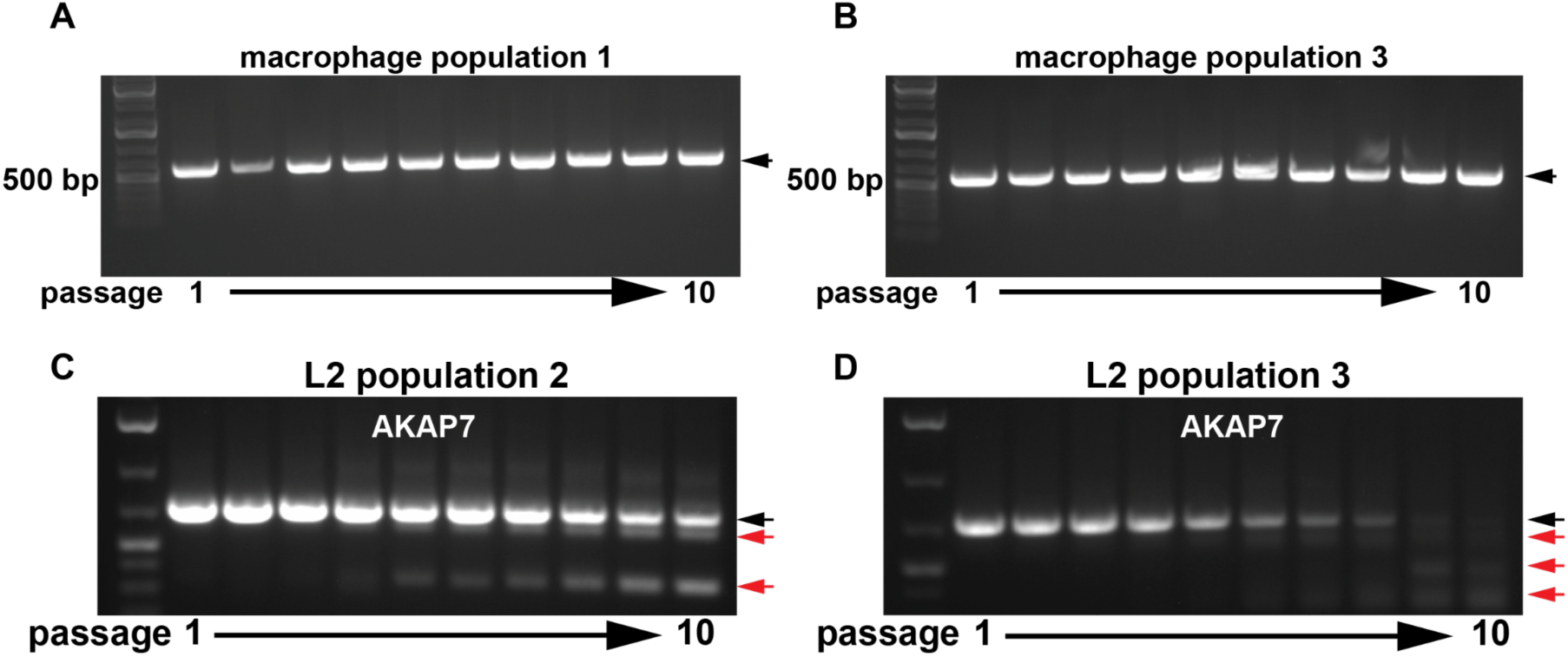
PCR analysis of AKAP7 during serial passage, related to Figure 1. A and B) AKAP7 PCR of passages 1 to 10 in macrophages. C and D) AKAP7 PCR of passages 1 to 10 in L2 experimental evolution replicate 2 and 3. Black arrows indicate full-length AKAP7, and red arrows indicate AKAP7 amplicons containing deletions.

**Figure S2.**
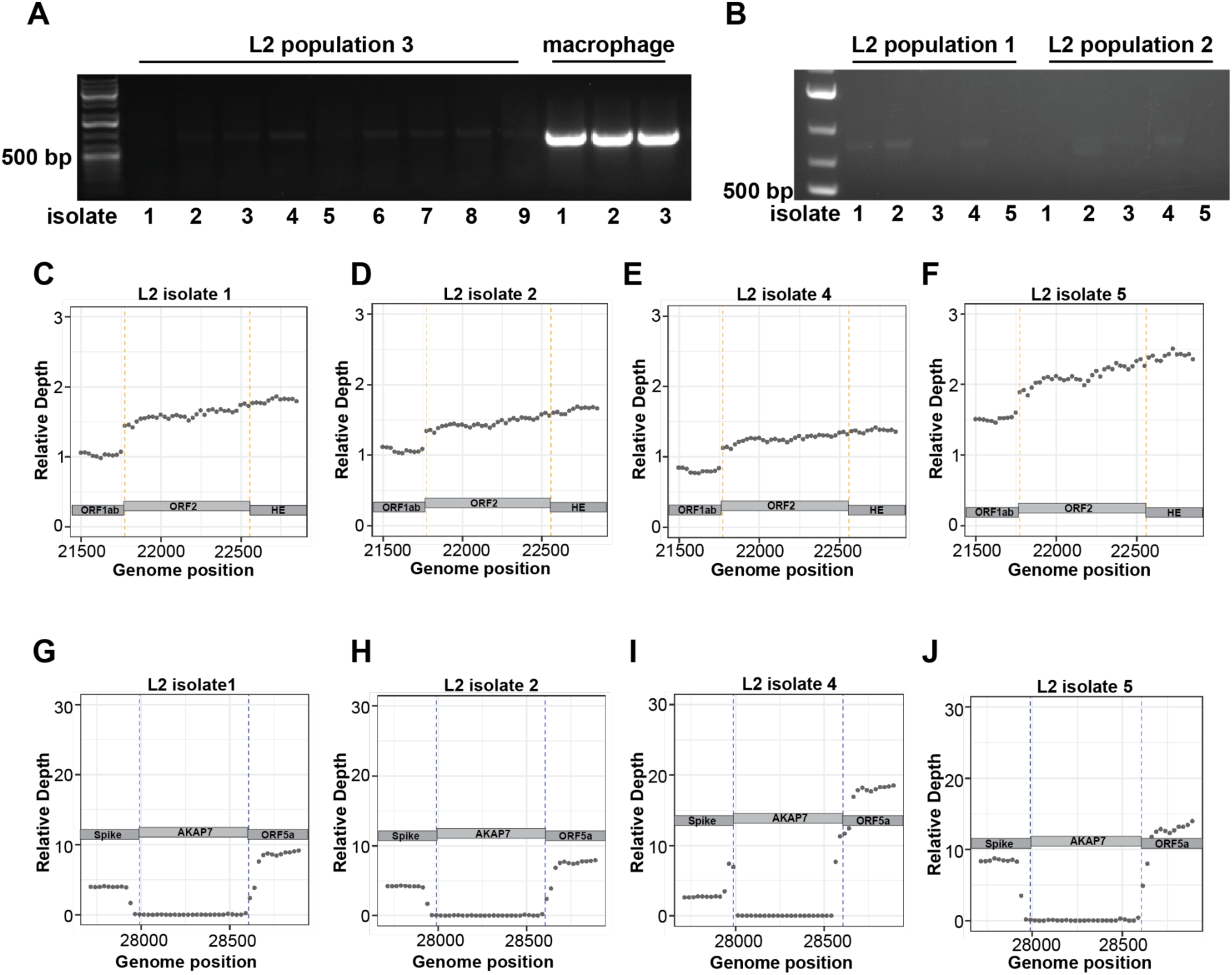
Direct cDNA sequencing shows loss of AKAP7 and retention of ORF2 during serial passage, related to Figure 2. A-B) PCR analysis of AKAP7 in plaque purified p10 MHV^AKAP7^ isolates. C-F) Relative coverage depth plots of ORF in purified plaque isolates from p10 L2 fibroblast purified plaque isolates. G-J) Relative coverage depth plots of AKAP7 in purified plaque isolates from p10 L2 fibroblast purified plaque isolates.

**Figure S3.**
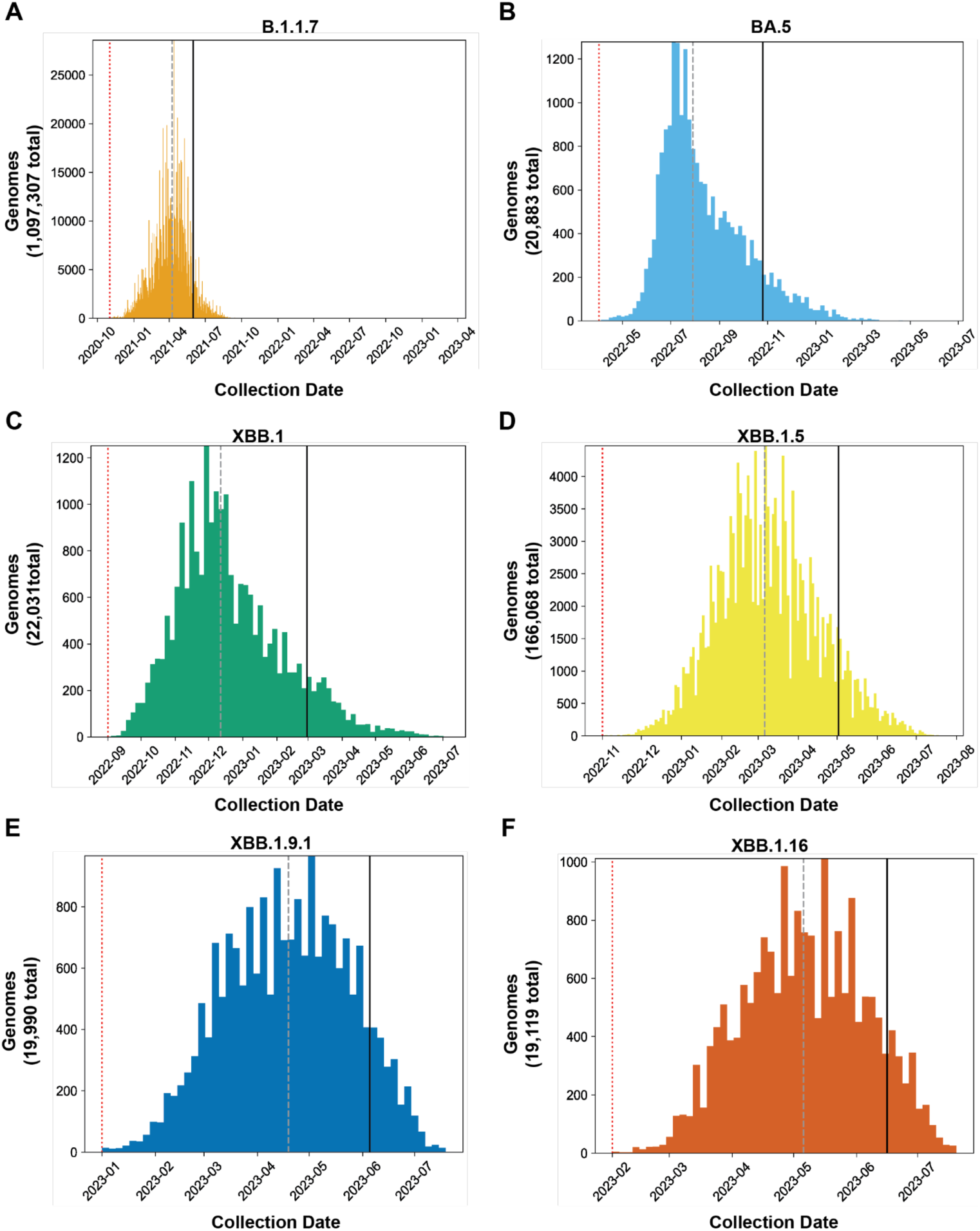
Histogram of ORF8 premature-stop codon-containing SARS-CoV-2 lineage genomes plotted by sampling date, related to Figures 4 and 5. A-E) Histograms of SARS-CoV-2 lineages we sampled that have premature stop codons in ORF8. The x-axis on each plot is the collection date and the y-axis indicates the number of genomes collected on each date. The red dotted line indicates the start-date of the lineage, determined by identifying the date when sequences reported from each lineage began to increase. The dashed line is the date by which 50% of all sequences assigned to this lineage were collected and the solid vertical line the date by 90% had been collected (as of 11/29/2023).

**Figure S4.**
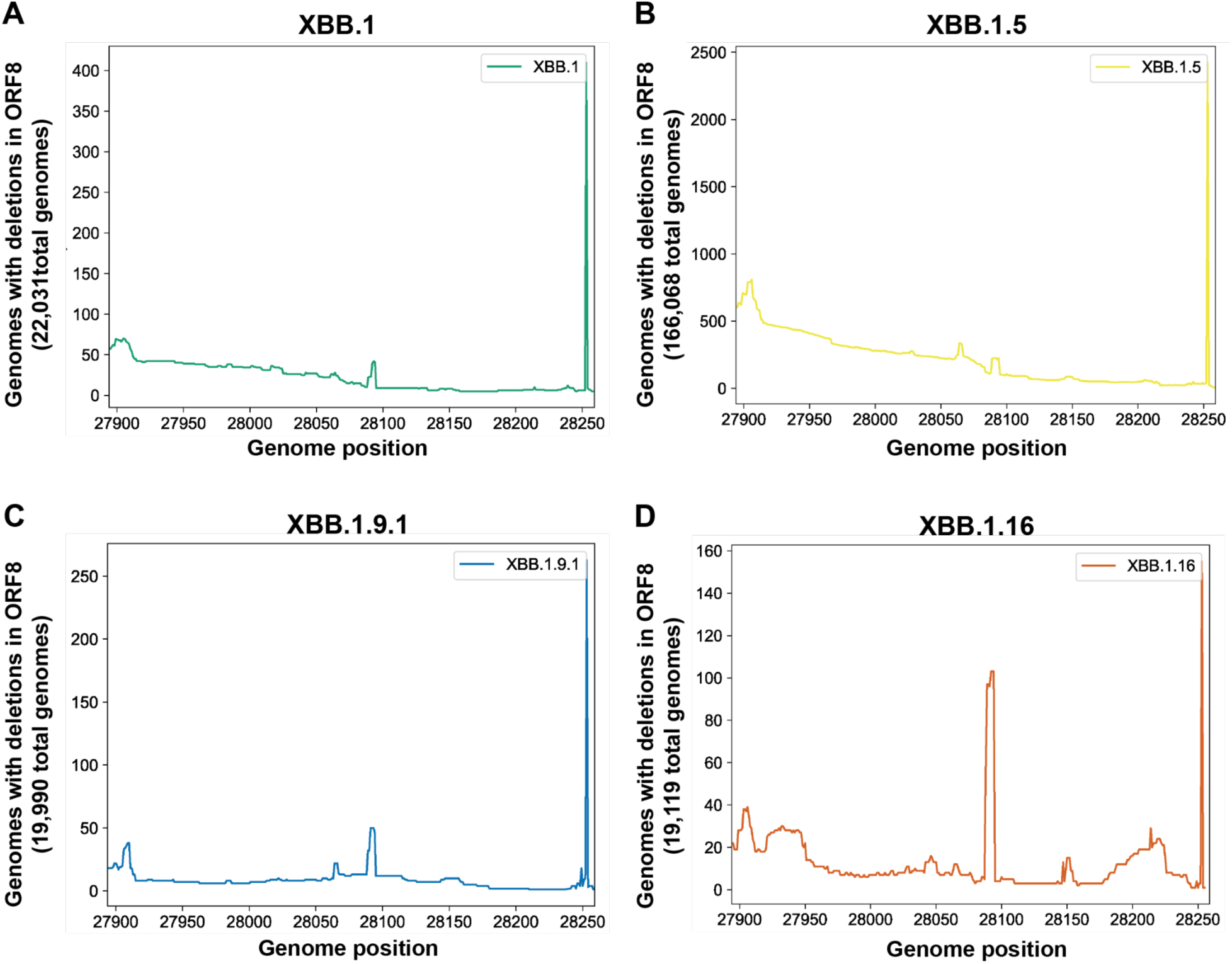
Deletions are not consistently enriched in discrete regions of SARS-CoV-2 ORF8, related to Figure 5. A-D) Plots depicting the number of sequences sampled on each date with deletions at each position within SARS-CoV-2 ORF8 for the indicated lineage.

**Table S3.** SARS-CoV-2 ORF8 deletion length distribution raw data, related to Figure 4C. This table contains raw data of deletion lengths in SARS-CoV-2 ORF8, specifically the percentage of deletions >10 nucleotides, 10-100 nucleotides, ad >100 nucleotides for all lineages in aggregate, and broken out into individual lineages.

|  | <10 nucleotides | 10-100 nucleotides | >100 nucleotides |
| --- | --- | --- | --- |
| **All lineages** | 97.38% | 1.94% | 0.68% |
| **B.1.1.7** | 98.16% | 1.36% | 0.49% |
| **BA.5** | 95.51% | 0.79% | 3.70% |
| **XBB.1** | 93.27% | 5.0% | 1.73% |
| **XBB.1.16** | 86.6% | 12.92% | 0.48% |
| **XBB.1.5** | 90.67% | 7.34% | 1.98% |
| **XBB.1.9.1** | 92.73% | 5.26% | 2.01% |

